# A rare missense mutation in *MYH6* confers high risk of coarctation of the aorta

**DOI:** 10.1101/180794

**Authors:** Thorsteinn Bjornsson, Rosa B. Thorolfsdottir, Gardar Sveinbjornsson, Patrick Sulem, Gudmundur L. Norddahl, Anna Helgadottir, Solveig Gretarsdottir, Audur Magnusdottir, Ragnar Danielsen, Emil L. Sigurdsson, Berglind Adalsteinsdottir, Sverrir I. Gunnarsson, Ingileif Jonsdottir, David O. Arnar, Hrodmar Helgason, Tomas Gudbjartsson, Daniel F. Gudbjartsson, Unnur Thorsteinsdottir, Hilma Holm, Kari Stefansson

## Abstract

Coarctation of the aorta (CoA) accounts for 4-8% of congenital heart defects (CHDs) and carries substantial morbidity despite treatment^1^. We performed a genome-wide association study (GWAS) of CoA among 120 Icelandic cases and 355,166 controls and found association with a rare (frequency = 0.34%) missense mutation p.Arg721Trp in *MYH6* (odds ratio (OR) = 44.2, *P* = 5.0x10^-22^), encoding an essential sarcomere protein. Approximately 20% of CoA cases in Iceland carry p.Arg721Trp. This is the first mutation associated with non-familial or sporadic CoA at a population level. P.Arg721Trp also associates with risk of bicuspid aortic valve (BAV) and other CHDs and has been reported to have a broad effect on cardiac electrical function and to associate strongly with sick sinus syndrome (SSS) and atrial fibrillation (AF)^2^. These findings suggest that p.Arg721Trp in *MYH6* causes a cardiac syndrome with highly variable expressivity, and emphasize the major importance of sarcomere integrity for cardiac development and function.

---

CoA is defined by local narrowing of the proximal descending aorta and/or aortic arch, causes left ventricular outflow tract obstruction (LVOTO), and typically presents as either heart failure in infants or hypertension later in life^3^. Surgical or interventional treatment considerably improves outcome but risk of premature cardiovascular morbidity and mortality remains despite appropriate therapy^4^.

CoA presents most commonly in a sporadic fashion and is accompanied by BAV in about half of cases, but less commonly with other CHDs^5^. Less frequently, it cosegregates in families with other LVOTO malformations including BAV, congenital aortic valve stenosis and hypoplastic left heart syndrome (HLHS)^6^. The LVOTO malformations as a group are markedly heritable (0.71-0.90) and have a high relative risk for first-degree relatives (36.9)^7^. In addition, around 15% of CoA cases occur as part of a recognized genetic syndrome (e.g. 45,X or Turner)^8^.

Not much is known about genetic causes of non-syndromic CoA. Several studies have found mutations in families with LVOTO malformations and a few cases of sporadic CoA, both with and without concomitant CHDs. The most strongly implicated gene is *NOTCH1*^9-11^, encoding a transmembrane receptor that regulates cell fate during development. Mutations in other genes, including *MYH6*^*12,13*^, *SMAD6*^14^, *NKX2-5*^15^ and *GATA5*^16^, have been found in one or few cases of CoA. The *MYH6* mutations were found in two families of CoA cases, one with predisposition to atrial septal defect (ASD)^12^ and the other to HLHS^13^. In addition, knockout animal models of several genes found within copy number variants in CoA cases in man, including *MCTP2*^*17*^, *MATR3*^*18*^ and *FOXC1*^*19*^, have been shown to result in CoA-like phenotypes.

To search for sequence variants that associate with non-syndromic CoA, we performed a GWAS including 120 Icelandic cases and 355,116 population controls. We tested 32.5 million sequence variants (SNPs and INDELs) identified through whole-genome sequencing (WGS) of 15,220 Icelanders (median coverage 32X) and imputed into 151,677 chip-typed, long-range phased Icelanders and their close relatives. The utilization of long-range phased haplotypes enables accurate imputation of variants with frequency down to approximately 0.02% in this data set. We performed association testing using logistic regression adjusting for gender, age and county. The threshold for genome-wide significance was set using a class-specific Bonferroni procedure based on functional impact of classes of variants^20^.

The 120 CoA cases were diagnosed between the years 1950 and 2016, with most cases (75%) diagnosed after 1990. We show the phenotypic characteristics of the CoA cases in **Supplementary Table 1**. About half were diagnosed during the first month of life and three quarters during the first year of life. As expected^4^, CoA was more common in males than in females (1.6:1). About three quarters of cases had other concomitant CHDs, most commonly BAV and ventricular septal defect (VSD). Similar to other studies^21^, the aortic valve was bicuspid in about half of CoA cases.

We observed a genome-wide significant association with CoA at chromosome 14q11 (Figure 1), explained by a rare (allele frequency = 0.34%) missense mutation p.Arg721Trp (c.2161C>T) in *MYH6*, encoding the alpha myosin heavy chain subunit (*α*MHC) in cardiac muscle. *α*MHC is a main component of the sarcomere, the basic contractile unit of cardiac muscle^22^. P.Arg721Trp associated with CoA with an OR of 44.2 (95% confidence interval (CI) 20.5 – 95.5) and *P* = 5.01 × 10^-22^ (genome-wide significance threshold for missense variants was set at 6.5 × 10^-8^, see Methods)^20^ (Figure 2). None of the genotyped cases or controls (N = 151,677) were homozygous for the mutation, consistent with its low frequency (1.8 homozygotes expected under Hardy-Weinberg equilibrium). Since we observed no homozygotes, we could not discriminate between the dominant and the multiplicative modes of inheritance.

**Figure 1.**
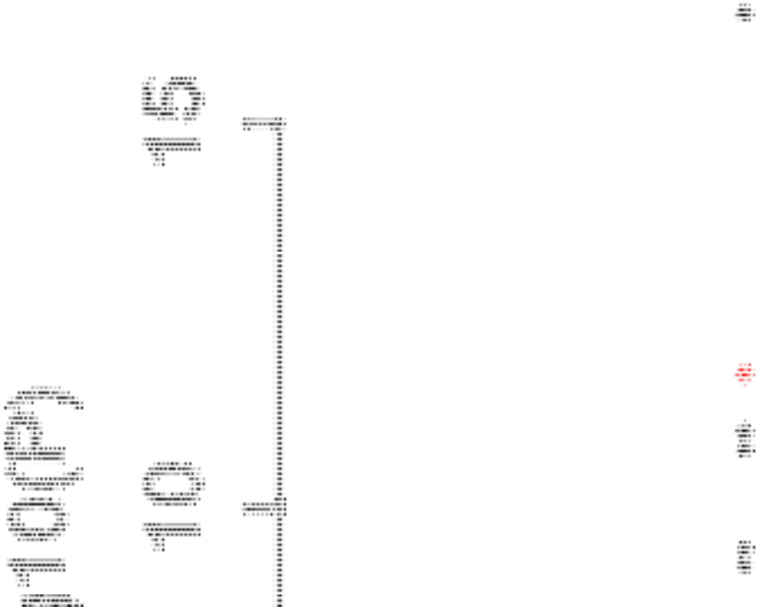
Manhattan plot of CoA genome-wide association study in Iceland. The *P* values (-log10) are plotted against their respective positions on each chromosome.

**Figure 2.**
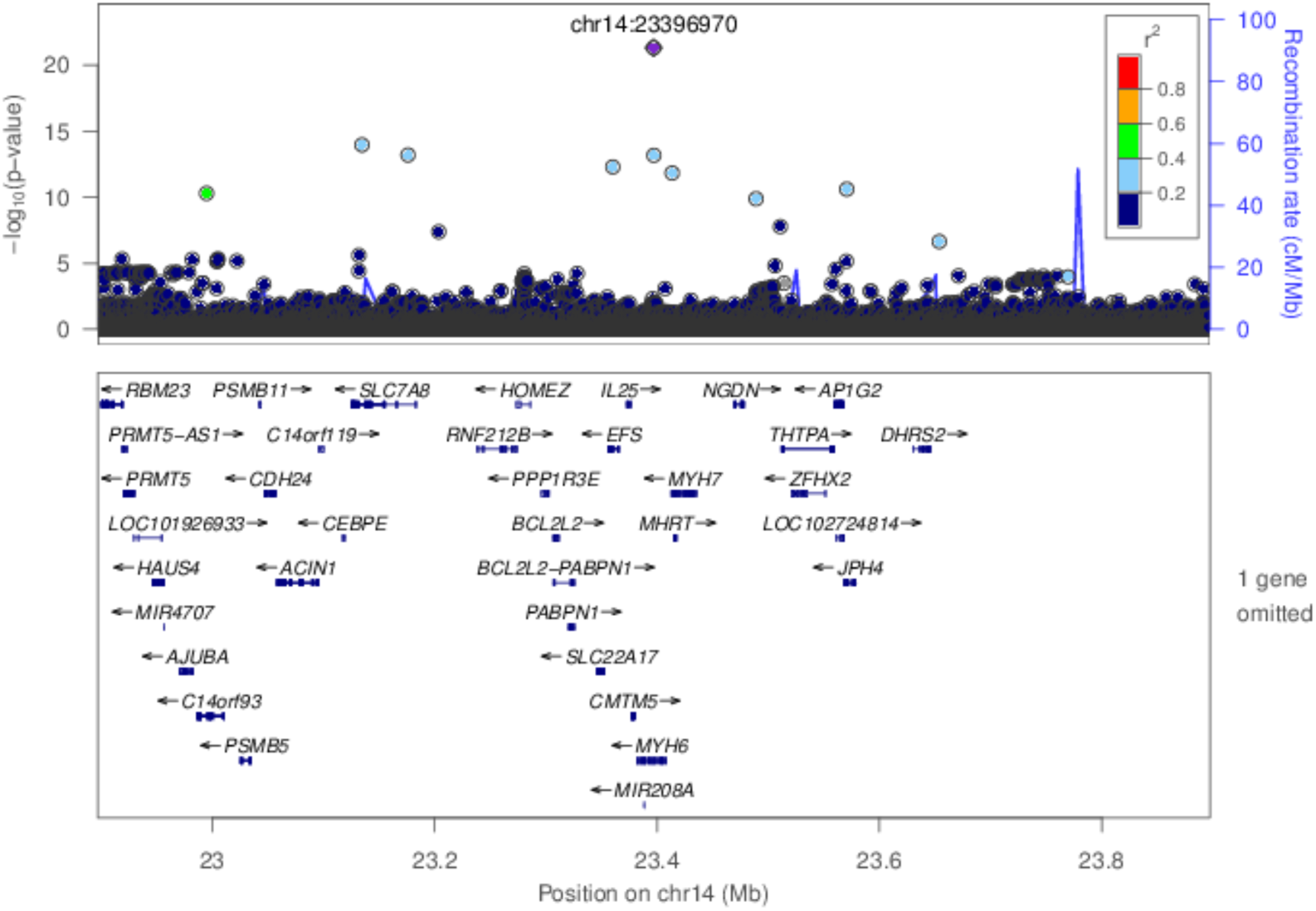
Region plot for the association of variants on 14q11 with CoA. Shown is a 1 Mb region on chromosome 14. The strongest association is with the missense variant p.Arg721Trp in *MYH6* located at position 23,396,970 on chromosome 14. The nine other variants shown are weakly correlated with p.Arg721Trp, *r*^2^ between 0.6-0.4 (green) and 0.4-0.2 (light blue).

The p.Arg721Trp mutation is located in exon 18 (out of 39 exons) of *MYH6* and leads to a missense arginine to tryptophan alteration at amino acid 721 (full-length protein 1939 amino acid) (**Supplementary Figure 1**). It is located in the converter domain of αMHC (**Supplementary Figure 1 and 2**), a small domain crucial in conveying a conformational change from the active site to the lever arm upon adenosine triphosphate (ATP) hydrolysis^23^. It is considered likely that the mutation alters protein function (SIFT = 0, PolyPhen = 0.99, MutationTaster = 0.93), probably altering the folding of the converter domain. There were 987 carriers of p.Arg721Trp among the 151,677 chip-typed Icelanders and eight of those (one per 123 carriers) were diagnosed with CoA. In line with low penetrance of the mutation for CoA, p.Arg721Trp carriers diagnosed with CoA did not cluster in families. However, 20% of the 39 chip-typed CoA cases carried p.Arg721Trp. Thus, while the penetrance of the mutation for CoA is low, it accounts for a large proportion of CoA cases in the Icelandic population.

To determine whether there were phenotypic differences between carriers (n=24) and non-carriers (n=96) of the p.Arg721Trp mutation within the CoA sample set, we evaluated the frequencies of various clinical characteristics between the two groups (**Supplementary Table 2**). Carriers were nominally more likely to present with mild rather than more critical and complex forms of CoA (OR = 4.2 and *P* = 0.023). We observed no other differences.

The p.Arg721Trp mutation is neither present in The Genome Aggregation Database (gnomAD), Cambridge; MA (http://gnomad.broadinstitute.org/) (October 2016) holding data from 126,216 exome sequences and 15,136 whole-genome sequenced unrelated individuals^24^, nor in the Exome Variant Server, containing sequence data from 6,503 individuals (Exome Variant Server, NHLBI Exome Sequencing Project (ESP), Seattle, WA, (http://evs.gs.washington.edu/EVS/) (v0029 2014; November). The p.Arg721Trp mutation thus appears to be absent from other populations or present at a very low frequency. The Icelandic population is a founder population in that a small number of ancestors account for a relatively large proportion of genetic diversity in the population. Hence, sequence variants that are very rare in more outbred populations may be more frequent in Icelanders, like p.Arg721Trp^25^.

Very rare mutations in *MYH6*, other than p.Arg721Trp, have been linked to various CHDs^12,13,26^, particularly familial ASD^27,28^ and both dilated and hypertrophic cardiomyopathy (HCM)^29^. In all instances, these mutations have been restricted to a few sporadic cases or to few families. In two of these families, one with predisposition to ASD^12^ and the other to HLHS^13^, some of the affected family members had other cardiac defects, including CoA. The *MYH6* p.Arg721Trp contrasts these rare familial mutations as it associates with CoA at the population level and explains about 20% of CoA cases in Iceland.

We have previously demonstrated that p.Arg721Trp associates strongly with SSS and AF, atrial arrhythmias that are common in the elderly and frequently coexist^2^ (Table 1). In the context of assessing effects of AF risk variants on cardiac conduction, we have also recently found that p.Arg721Trp associates with many ECG measures corresponding to a widespread effect on cardiac electrical function^30^ (**Supplementary Figure 3**).

**Table 1.**
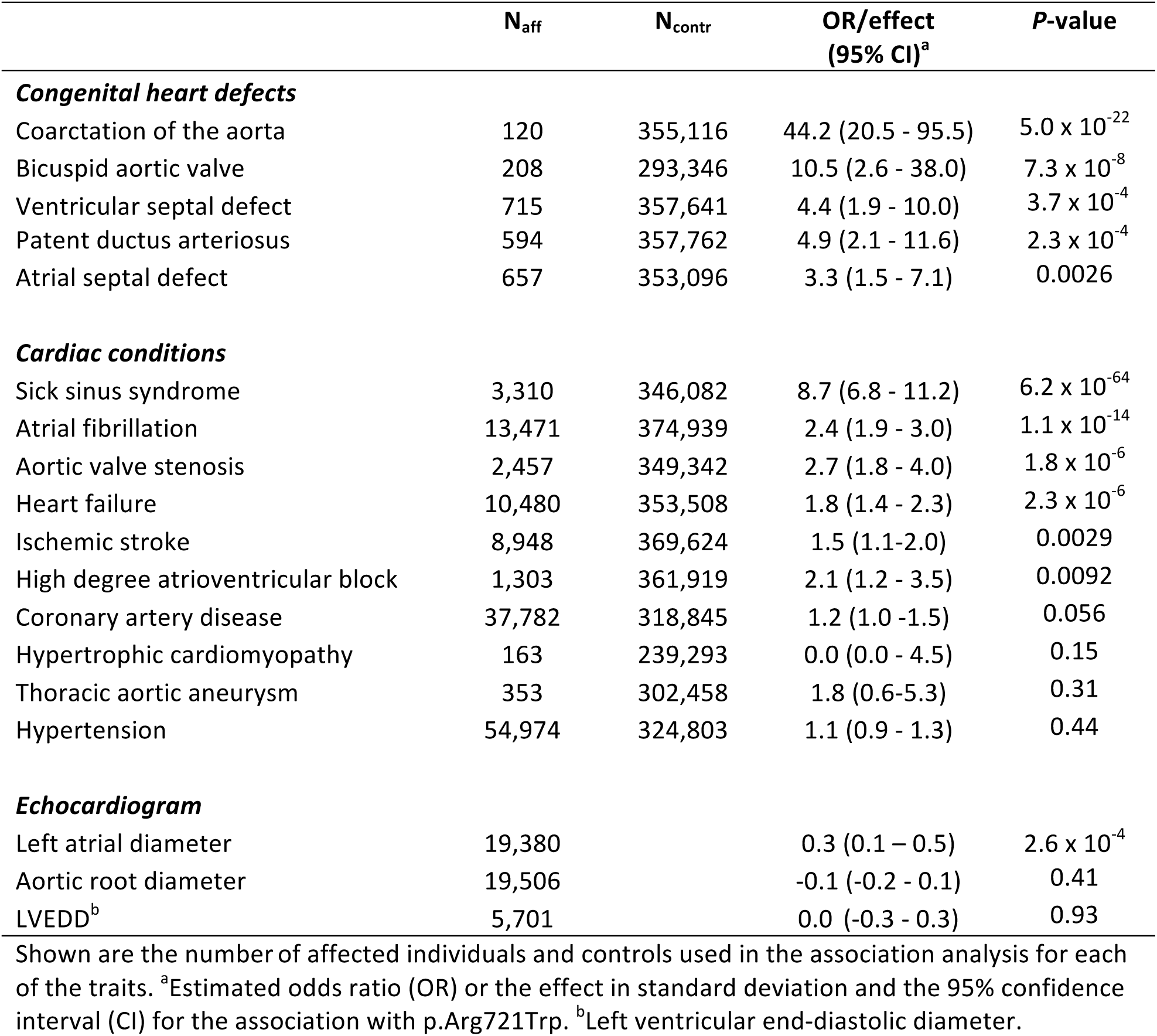
Association of p.Arg721Trp with congenital heart defects and various cardiac phenotypes.

|  | N<sub>aff</sub> | N<sub>contr</sub> | OR/effect<br>(95% CI)<sup>a</sup> | P-value |
| --- | --- | --- | --- | --- |
| ***Congenital heart defects*** |  |  |  |  |
| Coarctation of the aorta | 120 | 355,116 | 44.2 (20.5 - 95.5) | 5.0 x 10<sup>-22</sup> |
| Bicuspid aortic valve | 208 | 293,346 | 10.5 (2.6 - 38.0) | 7.3 x 10<sup>-8</sup> |
| Ventricular septal defect | 715 | 357,641 | 4.4 (1.9 - 10.0) | 3.7 x 10<sup>-4</sup> |
| Patent ductus arteriosus | 594 | 357,762 | 4.9 (2.1 - 11.6) | 2.3 x 10<sup>-4</sup> |
| Atrial septal defect | 657 | 353,096 | 3.3 (1.5 - 7.1) | 0.0026 |
| ***Cardiac conditions*** |  |  |  |  |
| Sick sinus syndrome | 3,310 | 346,082 | 8.7 (6.8 - 11.2) | 6.2 x 10<sup>-64</sup> |
| Atrial fibrillation | 13,471 | 374,939 | 2.4 (1.9 - 3.0) | 1.1 x 10<sup>-14</sup> |
| Aortic valve stenosis | 2,457 | 349,342 | 2.7 (1.8 - 4.0) | 1.8 x 10<sup>-6</sup> |
| Heart failure | 10,480 | 353,508 | 1.8 (1.4 - 2.3) | 2.3 x 10<sup>-6</sup> |
| Ischemic stroke | 8,948 | 369,624 | 1.5 (1.1-2.0) | 0.0029 |
| High degree atrioventricular block | 1,303 | 361,919 | 2.1 (1.2 - 3.5) | 0.0092 |
| Coronary artery disease | 37,782 | 318,845 | 1.2 (1.0 -1.5) | 0.056 |
| Hypertrophic cardiomyopathy | 163 | 239,293 | 0.0 (0.0 - 4.5) | 0.15 |
| Thoracic aortic aneurysm | 353 | 302,458 | 1.8 (0.6-5.3) | 0.31 |
| Hypertension | 54,974 | 324,803 | 1.1 (0.9 - 1.3) | 0.44 |
| ***Echocardiogram*** |  |  |  |  |
| Left atrial diameter | 19,380 |  | 0.3 (0.1 – 0.5) | 2.6 x 10<sup>-4</sup> |
| Aortic root diameter | 19,506 |  | -0.1 (-0.2 - 0.1) | 0.41 |
| LVEDD<sup>b</sup> | 5,701 |  | 0.0 (-0.3 - 0.3) | 0.93 |

Shown are the number of affected individuals and controls used in the association analysis for each of the traits. <sup>a</sup>Estimated odds ratio (OR) or the effect in standard deviation and the 95% confidence interval (CI) for the association with p.Arg721Trp. <sup>b</sup>Left ventricular end-diastolic diameter.

To further explore the effect of the p.Arg721Trp *MYH6* mutation, we tested it for association with additional cardiac phenotypes, including other CHDs, common heart diseases and several echocardiogram variables (Table 1, **Supplementary Table 3 and Supplementary Figure 3;** significance threshold set at P < 0.003 (0.05 / 17 individual phenotypes tested). The p.Arg721Trp mutation associated with increased risk of several CHDs: BAV, VSD, ASD and patent ductus arteriosus (Table 1). As expected, the strongest association was with BAV (OR = 10.5 and *P* = 7.3 × 10^-8^). In addition, the mutation associated with late onset aortic valve stenosis. To assess if p.Arg721Trp associates with CHDs in the absence of diagnosed CoA, we re-tested for association after removing CoA cases from the analysis. Although the effect of p.Arg721Trp is consistently weaker, the associations remain (**Supplementary Table 3**). We cannot exclude the existence of undiagnosed CoA in these cases. The mutation also associated with heart failure and ischemic stroke and with diameter of the left atrium but not with other variables derived from the echocardiographic data such as aortic root diameter or left ventricular end-diastolic diameter (Table 1). The p.Arg721Trp mutation did not associate with hypertension or coronary artery disease.

The major components of cardiac muscle myosin are *α*MHC and beta myosin heavy chain (βMHC) encoded by *MYH6* and *MYH7*, respectively. Both *α*MHC and βMHC are expressed throughout the heart during embryonic cardiogenesis whereas in the adult heart *α*MHC is predominantly expressed in the atrium^31^. Expression of *MYH6* has not been detected in the aorta (http://gtexportal.org).

The pathogenesis of CoA is not well understood. One of the main models of CoA pathogenesis, the hemodynamic theory^3,32^, stipulates that cardiac lesions resulting in decreased left ventricular outflow promote development of CoA by reducing blood flow through the fetal aorta. The p.Arg721Trp mutation could predispose to CoA by reducing blood flow through the fetal aorta because of diminished contraction of the developing heart. This hypothesis is supported by overexpression studies in rat cardiomyocytes showing that the p.Arg721Trp mutation impairs sarcomeric structure^33^ and by our ECG data demonstrating widespread effect of p.Arg721Trp on cardiac electrical function, including in the ventricles. Our hypothesis is compatible with the fact that *MYH6* is expressed in the ventricles during the development of the heart but not in the aorta.

In summary, through GWAS based on WGS, we found a rare missense mutation in the sarcomere gene *MYH6* that has a strong effect on the risk of CoA in the Icelandic population and explains a substantial fraction of Icelandic CoA cases. The same mutation also associates with other CHDs, in particular BAV. It has a widespread effect on cardiac electrical function and associates strongly with atrial arrhythmias, both SSS and AF, suggesting a cardiac syndrome with highly variable expressivity that is difficult to identify clinically without sequence information. This is the first mutation associated with non-familial or sporadic form of CoA at a population level. This discovery gives important insights into the pathophysiology of CoA, supporting the hemodynamic theory of defect development.

## Online methods

### Icelandic coarctation of the aorta study population

The coarctation of the aorta (CoA) sample set included Icelanders who received the discharge diagnosis of CoA at Landspitali, The National University Hospital (LUH) in Reykjavik, the only tertiary referral center in Iceland, between 1984 and 2016. LUH is the only hospital in Iceland with specialized care for patients with congenital heart defects (CHDs). CoA cases were identified either through diagnosis codes of CoA (ICD-9 code 747.1, ICD-10 code Q25.1) registered between 1990 and 2016 or procedure codes of CoA (WHO codes 1-273, 5-369, 5-382 and 5-387, NOMESCO codes FDJ 00, FDJ 10, FDJ 20, FDJ 30, FDJ 42 and FDJ 96) registered between 1984 and 2016. CoA was defined as a congenital narrowing of aorta, the diagnosis of which was confirmed by echocardiography and/or cardiac catheterization by a cardiologist. The diagnoses of CoA and other CHDs were confirmed and type of CoA established through review of electronic and paper medical records at LUH. We identified 146 Icelanders with CoA, 11 of which were syndromic and excluded from the study. Of the remaining 135 cases, genotypes were available for 120 cases that were used in the analysis. The controls used in the CoA case-control analyses of this study consisted of disease-free controls randomly drawn from the Icelandic genealogical database and individuals from other genetic studies at deCODE. The National Bioethics Committee of Iceland approved the study.

Based on severity and/or anatomy of the narrowing, we classified the CoA cases into five different types. Mild CoA was defined as untreated, hemodynamically insignificant CoA. Moderate CoA was defined as treated, hemodynamically significant CoA. Critical CoA was defined as severe narrowing presenting as either congestive heart failure or cardiogenic shock during the neonatal period, with or without concomitant large ventricular septal defect/s (VSD). Complex CHD was defined as CoA occurring in conjunction with multiple CHDs. Finally, we included coarctation of the abdominal aorta and interruption of the aortic arch type A. We also documented the age at diagnosis and the presence of associated hypoplasia, CHDs and *MYH6* related disease. Information on date of birth and gender came from the Book of Icelanders (contains genealogical information about Icelanders). The phenotypic information was used for phenotype-genotype correlation of CoA cases.

### The deCODE genetics phenotype database

The deCODE genetics phenotype database contains extensive medical information on various diseases and traits. The CHD sample sets included 715 individuals with ventricular septal defect (VSD) (ICD 10: Q21.0 or ICD 9: 745.4), 657 individuals with atrial septal defect (ASD) (ICD 10: Q21.1 or ICD 9: 745.5), 594 individuals with patent ductus arteriosus (ICD 10: Q25.0 or ICD 9: 747.0) and 208 individuals with description of bicuspid aortic valve (BAV) according to cardiologist interpretation of echocardiograms from LUH between 1994 and 2015. The coronary artery disease dataset (N=37,782) has been described^34^. The aortic valve stenosis (N=2,457), atrial fibrillation (N=13,471), heart failure (N=10,480), high-degree atrioventricular block (N=1,303), ischemic stroke (N=8,948), sick sinus syndrome (N=3,310) and thoracic aortic aneurysm (N=353) sample sets were based on discharge diagnoses from LUH from 1987 to 2015. The hypertension sample set included 54,974 individuals who received the diagnosis of hypertension at LUH or the Primary Health Care Clinics of the Reykjavik area. Patients with a clinical diagnosis of hypertrophic cardiomyopathy (HCM) were identified from surveys of electronic medical records and a review of Icelandic echocardiogram databases, including LUH, Akureyri Hospital, and private cardiology clinics^35^. In total, 163 Icelandic patients with available genotype information fulfilled the following clinical criteria for HCM: LV maximal wall thickness ≥15 mm in the absence of another cardiac or systemic disease capable of producing a similar magnitude of hypertrophy.

Three phenotypes describing measurements from echocardiograms were used in the analysis, left atrial diameter (N = 19,380), aortic root diameter (19,506) and left ventricular end-diastolic diameter (N = 5,701). These measurements were obtained from a database of 53,122 echocardiograms from 27,460 individuals performed and documented by a cardiologist at LUH between 1994 and 2015. Measurements were adjusted for sex, year of birth and age at measurement and were subsequently standardized to have a normal distribution. Electrocardiogram (ECG) data was collected from LUH in Reykjavik and included all ECGs obtained and digitally stored from 1998 to 2015, a total of 434,000 ECGs from up to 88,217 individuals.

### Genotyping, imputation and whole-genome sequencing

For chip genotyping, 151,677 samples were typed with the Illumina HumanHap300, HumanCNV370, HumanHap610, HumanHap1M, HumanHap660, Omni-1, Omni 2.5 or Omni Express bead chips at deCODE genetics. Chip SNPs were excluded if they had (1) yield less than 95%, (2) minor allele frequency less than 1% in the population or (3) significant deviation from Hardy-Weinberg equilibrium (*P*<0.001), (4) if they produced an excessive inheritance error rate (over 0.001) and (5) if there was substantial difference in allele frequency between chip types (from just a single chip if that resolved all differences, but from all chips otherwise). All samples with a call rate below 97% were excluded from the analysis. The final SNP set used for long-range phasing comprised 676,913 autosomal SNPs. Long range phasing of all chip-genotyped individuals was performed with methods described previously^36,37^. In brief, phasing is achieved by using an iterative algorithm that phases a single proband at a time given the available phasing information about everyone else that shares a long haplotype identically by state with the proband. Given the large fraction of the Icelandic population that has been chip-typed, accurate long range phasing is available genome-wide for all chip-typed Icelanders. For long-range phased haplotype association analysis, we then partitioned the genome into non-overlapping fixed 0.3 cM bins. Within each bin, we observed the haplotype diversity described by the combination of all chip-typed markers in the bin.

The whole genomes of 15,220 Icelanders were sequenced using Illumina technology to a median depth of 32X. This dataset contains samples obtained using three different library preparation methods from Illumina. In addition, sequencing was performed using three different types of Illumina sequencing instruments. (A) Standard TruSeq DNA library preparation method using Illumina GAIIx and/or HiSeq 2000 sequencers; (B)TruSeq DNA PCR-free library preparation method using Illumina HiSeq 2500 sequencers and (C) TruSeq Nano DNA library preparation method using Illumina HiSeq X sequencers.

In the sequencing dataset SNPs and INDELs were identified and genotypes called using joint calling with the Genome Analysis Toolkit HaplotypeCaller (GATK version 3.3.0)^38^. Genotype calls were improved by using information about haplotype sharing, taking advantage of the fact that all the sequenced individuals had also been chip-typed and long-range phased.

The sequence variants identified in the 15,220 sequenced Icelanders were then imputed into 151,677 Icelanders who had been genotyped with various Illumina SNP chips and their genotypes phased using long-range phasing^36,37^. The imputation into the chip typed long-range phased individuals was performed with the same model as used by IMPUTE^39^. Using genealogic information on Icelanders from The Book of Icelanders^40^, the sequence variants were imputed into first and second-degree relatives of array genotyped individuals to further increase the sample size for association analysis and to increase the power to detect associations. We identified 32.5 million high quality sequence variants (all with imputation information >0.8 that mapped to build hg38) that were tested for association with CoA under the multiplicative model.

### Association analysis

Association testing for case-control analysis was performed using logistic regression, adjusting for gender, age and county. A total of 32.5 million variants were used in the association analysis under a multiplicative model.

To account for inflation in test statistics due to cryptic relatedness and stratification, we applied the method of LD score regression^41^. With a set of 1.1M variants, we regressed the χ^2^ statistics from our genome-wide association scan against LD score and used the intercept as a correction factor. The LD scores were downloaded from a LD score database^41^. The estimated correction factor was 1.04 for the multiplicative model of the CoA association.

To correct for multiple testing we used the weighted Holm-Bonferroni method^42^ to allocate family wise error rate of 0.05 equally between four annotation-based classes of sequence variants. For the multiplicative model this yielded significance thresholds of 2.6 × 10^-7^ for high-impact variants (including stop-gained, frameshift, splice acceptor or donor, N = 8,474), 5.1 × 10^-8^ for moderate-impact variants (including missense, splice-region variants and in-frame INDELs, N = 149,983), 4.6 × 10^-9^ for low impact variants (including synonymous variants 3’ and 5’ UTR variants, N = 2,283,889), 2.3 × 10^-9^ for intergenic and deep intronic variants overlapping DNase hypersensitive sites (N = 3,913,058) and other variants 7.9 × 10^-10^ (intergenic and deep intronic, N = 26,108,039)^20^.

When testing the association of p.Arg721Trp with several other cardiac phenotypes, the controls used consisted of disease-free controls randomly drawn from other genetic studies at deCODE.

### Phenotypic differences between carriers and non-carriers of p.Arg721Trp

To analyze if CoA carriers of the p.Arg721Trp mutation differed clinically from non-carrier CoA cases, we evaluated the frequencies of various clinical characteristics in these two groups of CoA cases (**Supplementary Table 1**). Fisher’s exact test was used to test for significant difference in the mean frequency of the variants between non-carriers and carriers, and the odds ratio (OR) was calculated as (pa/(1-pa)) / (pc/(1-pc)), where pa and pc are the mean frequencies of the variants in non-carriers and carriers, respectively.

## Acknowledgements

We thank all the study subjects for their valuable participation as well as our colleagues that contributed to data collection, sample handling and genotyping.

## Author information

### Affiliations

**deCODE genetics/Amgen, Inc., Reykjavik, Iceland.**

Thorsteinn Bjornsson, Rosa B. Thorolfsdottir, Gardar Sveinbjornsson, Patrick Sulem, Gudmundur L. Norddahl, Anna Helgadottir, Solveig Gretarsdottir, Audur Magnusdottir, Ingileif Jonsdottir, Daniel F. Gudbjartsson, Unnur Thorsteinsdottir, Hilma Holm, Kari Stefansson

**Department of Medicine, Landspitali – The National University Hospital of Iceland, Reykjavik, Iceland.**

Ragnar Danielsen, David O. Arnar

**Department of Family Medicine, University of Iceland, Reykjavik, Iceland.**

Emil L. Sigurdsson

**Department of Development, Primary Health Care of the Capital Area, Reykjavik, Iceland.**

Emil L. Sigurdsson

**Department of Cardiology, Haukeland University Hospital, Bergen, Norway.**

Berglind Adalsteinsdottir

**Faculty of Medicine, University of Iceland, Reykjavik, Iceland.**

Berglind Adalsteinsdottir, Ingileif Jonsdottir, David O. Arnar, Tomas Gudbjartsson, Unnur Thorsteinsdottir, Kari Stefansson

**Division of Cardiovascular Medicine, Department of Medicine, University of Wisconsin, Madison, WI, USA.**

Sverrir I. Gunnarsson

**Department of Immunology, Landspitali, the National University Hospital of Iceland, Reykjavik, Iceland.**

Ingileif Jonsdottir

**Children’s Hospital, Landspitali – The National University Hospital of Iceland, Reykjavik, Iceland.**

Hrodmar Helgason

**Department of Cardiothoracic Surgery, Landspitali – The National University Hospital of Iceland, Reykjavik, Iceland.**

Tomas Gudbjartsson

**School of Engineering and Natural Sciences, University of Iceland, Reykjavik, Iceland.**

Daniel F. Gudbjartsson

## Contributions

T.B., R.B.T., G.S., D.F.G., U.T., H. Holm and K.S. designed the study and interpreted the results. D.F.G. and G.S. performed the bioinformatics analysis, imputation and association analysis in the data sets. T.B., R.B.T., G.S., P.S., G.L.N., A.M., D.F.G., U.T. and H. Holm contributed to analysis of the data. T.B., R.B.T., A.H., S.G., R.D., E.L.S., B.A., S.I.G., I.J., D.O.A., H. Helgason, T.G. and H. Holm were responsible for phenotype data acquisition. T.B., R.B.T., G.S., D.F.G., U.T. and H. Holm wrote the initial draft. All authors reviewed and approved the final manuscript. K.S. supervised the study.

## Corresponding authors

Correspondence to: Kari Stefansson or Hilma Holm

